# Eg5 activity and density-driven bundling organize the human metaphase mitotic spindle independently of spindle bipolarity

**DOI:** 10.64898/2026.01.16.699769

**Authors:** William Conway, Daija Bobe, Vitaly Zimyanin, Gunar Fabig, Alex de Marco, Stefanie Redemann

## Abstract

The mitotic spindle segregates chromosomes through the coordinated actions of microtubules and molecular motors. Classic models propose that microtubules nucleate at spindle poles and grow inward to capture chromosomes; however, recent structural studies show that spindles contain short microtubules that do not span the distance between poles and chromosomes. It is unclear how these short microtubules assemble a bipolar spindle. Using cryo-electron tomography to map microtubule polarity in human metaphase spindles, we find that microtubules form locally antiparallel bundles with consistent 8 nm wall-to-wall spacing. We utilized motor perturbations and centriole depletion, which generated motor-active monopolar spindles, to reveal that the kinesin-5 motor Eg5 organizes local antiparallel overlap independently of spindle bipolarity. We found that bundles are organized by density-driven steric interactions rather than motor-mediated crosslinking. These findings support a self-organized, bottom-up model in which local microtubule-motor interactions within dense bundles generate forces that build the bipolar spindle, challenging pole-centric models.

**KEY FINDINGS:**

- Microtubules in metaphase spindles organize in locally antiparallel bundles with consistent 8 nm wall-to-wall spacing.
- The Eg5 motor generates antiparallel microtubule overlap independently of spindle bipolarity.
- Microtubule spacing scales with density through steric interactions, not direct motor crosslinking.
- Dynein regulates microtubule density to control bundle architecture; balanced Eg5-dynein density regulation maintains spindle bipolarity.

## INTRODUCTION

Cell division is a fundamental life process whose defects can cause cancer^1,2^ and lead to infertility^3,4^. During cell division, the segregation of chromosomes is driven by the spindle, a self-organized, ellipsoidal, bipolar structure composed of microtubules and numerous associated proteins^5–8^. Constituent microtubules are polar filaments that rapidly polymerize from minus-end to plus-end, depolymerize within seconds, and continually slide relative to each other^9–11^. The spindle is shaped by interactions of these microtubules with molecular motors, which walk along the surface of microtubules towards either the plus- or minus-end^12–15^. Motors can crosslink^16^, slide^11^, or transport microtubules^8,17^ throughout the spindle. In human spindles, the two most prominent motors are the plus-end-directed kinesin-5, Eg5^16,18^, and the minus-end-directed dynein^6,19^. Inhibition of either motor reshapes the spindle: inhibiting Eg5 collapses the spindle poles into an aster^20–23^, while inhibiting dynein defocuses the spindle poles to form a disordered, web-like structure^24–27^. Remarkably, inhibiting both motors restores the wild-type bipolar spindle structure, allowing cells to proceed into anaphase and segregate chromosomes^6,19^.

Traditional models of spindle assembly propose that microtubules nucleate at centrioles located at the spindle poles and grow inwards to search for, align, and segregate chromosomes in the spindle bulk^28–30^. A growing body of evidence suggests that this pole-centric view of spindle assembly is incomplete. Bipolar spindles can still form in the absence of centrioles and proceed to segregate chromosomes in anaphase^31–33^. *In vitro*, microtubules can nucleate around artificial chromosome beads and assemble a spindle in the absence of spindle poles^8,34,35^. Microtubules can also nucleate from existing microtubules using branching factors such as Augmin, providing additional routes for centriole-independent microtubule assembly^36–40^. An understanding of how these spindles assemble and segregate chromosomes in the absence of centriolar anchors, or equivalent MTOC-organized spindle poles observed in meiotic spindles and plant cells, has remained elusive.

Though dynamic data from light microscopy and *in vitro* reconstitution of spindle components have been very informative for investigating how spindles assemble, these approaches do not provide sufficient resolution to study the individual microtubule and molecular motor building blocks in their native context. To overcome this limitation, we^41–44,61^, along with several other groups^45–51^, have previously employed serial-section resin-embedded electron tomography, which achieves high enough resolution to visualize individual microtubules in the dense spindle. From these reconstructions, we have discovered that the spindle is composed of relatively short (∼2 µm) microtubules that form a connected network between the poles, in contrast to the traditional picture of long continuous microtubules originating at the poles and searching for chromosomes in the spindle midzone.

It is unclear how these short, interconnected microtubules interact with molecular motors to assemble the spindle and segregate chromosomes. Though plastic spindle reconstructions can visualize individual microtubules, resolution is compromised by bulky, heavy metal stains used to visualize proteins in the resin^52,53^. As a result, the polarity of individual microtubules and the locations of individual molecular motors, critical to developing a mechanistic picture of spindle assembly, cannot be reliably determined directly from imaging in plastic reconstructions^54^.

Here, we employ recent advances in FIB milling^55–57^ and cellular cryo-electron tomography^53,58,59^ (cryoET) to obtain near-atomic-resolution images of mitotic RPE-1 cells, enabling us to visualize the polarity of individual microtubules in the spindle. In contrast to predictions from a pole-centric model, we found that spindles are composed of locally antiparallel microtubule bundles with a consistent 8 nm wall-to-wall spacing between adjacent microtubules. By perturbing the two key molecular motors, Eg5 and dynein, as well as centriole duplication, we found that the Eg5 motor organizes antiparallel microtubule interactions independently of spindle bipolarity. Dynein is responsible for regulating microtubule density, which in turn sets the spacing between microtubules in dense bundles, crucial to maintaining spindle bipolarity. Taken together, these results favor a self-organized, bottom-up, mechanistic picture of spindle assembly, in which local interactions between microtubules and motors within dense bundles generate forces that assemble a bipolar spindle.

## RESULTS

### Bulk microtubule polarity in the spindle is not consistent with a model where microtubules are nucleated from the poles or by branching nucleation

In recent years, several intermediate-resolution structures of complete spindles have been generated by large-scale 3D electron tomography of high-pressure-frozen, resin-embedded samples^41–43,45,60,61^. We first examined a previously generated reconstruction of three human metaphase mitotic spindles in HeLa cells^42,43^. These reconstructed spindles are composed of short (mean length ∼2 µm) microtubules with a majority of the microtubules (∼60±5%) located more than 2 µm from either pole, suggesting that these microtubules are not centrosome-nucleated (Figure 1A). Very few centriolar microtubules are long enough to cross the metaphase plate (Figure 1B, C, Figure 1 – supplement 1). It is not clear how these short microtubules, which do not cross the metaphase plate, can establish a connection between the two spindle halves required to maintain a bipolar spindle.

**Figure 1:**
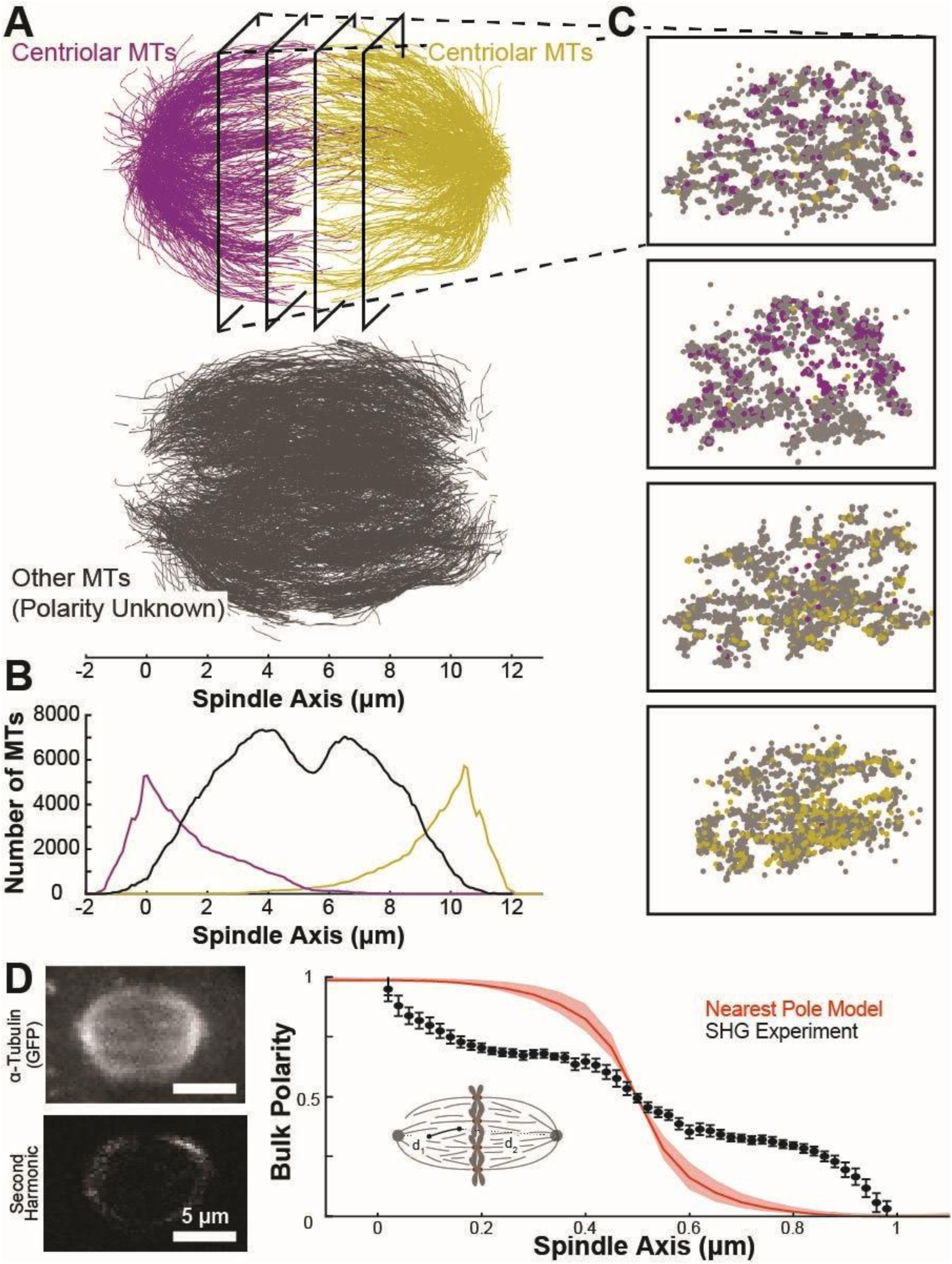
A model assigning microtubule polarity by the nearest pole cannot explain the spatial distribution of spindle bulk polarity. A) Reconstruction of microtubule trajectories from serial-section, resin-embedded electron tomography of a metaphase HeLa spindle. Microtubules are categorized by distance from either pole. Top: microtubules within 2 µm of the left (purple) or right (yellow) pole. Bottom: microtubules >2 µm from either pole (gray). B) Density of microtubules within 2 µm of the left pole (purple), within 2µm of the right pole (yellow), and more than 2 µm away from either pole (gray). C) Cross-sections of microtubule reconstructions taken orthogonal to the spindle axis at (-3 µm, -1 µm, 1 µm, 3 µm) from the center of the two poles. D) Comparison of bulk polarity measured by second harmonic with a model where the polarity of each microtubule in the plastic reconstructions is assigned to the nearest spindle pole. Left: Representative simultaneous images of GFP::α-tubulin and second harmonic signal in metaphase HeLa cells. Right: Bulk polarity in the spindle predicted by the nearest-pole-model (red) and calculated from the second harmonic data binned along the spindle axis (black), 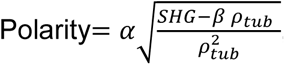, where 𝜌_𝑡𝑢𝑏_ is the density of tubulin, SHG is the second harmonic intensity, and the 𝛼 and 𝛽 constants are set by the spindle boundary conditions as previously described and validated^65^. N=14 cells, Error Bars: Standard error of the mean.

In previous work in budding yeast spindles, microtubule polarity was inferred from intermediate-resolution plastic reconstructions because all the microtubules originate at the spindle poles^62^; however, polarity cannot be determined directly from imaging in plastic due to limited resolution^52–54^. Inspired by this, and the suggestion that the non-centriolar microtubules are primarily nucleated by branching from existing microtubules^8,36,37,40^, we initially assigned the polarity of each microtubule to the nearest pole (Figure 1D, inset). To investigate whether this approach was reasonable, we used light microscopy (second harmonic generation, SHG) to quantify the bulk polarity of microtubules in HeLa spindles at fluorescent resolution (Figure 1D). This approach has been previously used to determine polarity in *C. elegans* and *Xenopus* egg extract spindles and has been validated by measuring microtubule polarity from depolymerization waves triggered by laser ablation experiments^17,63–65^. Using this approach, we observed that the polarity of microtubules is almost completely parallel at the poles and then falls off and plateaus in the spindle bulk, and then becomes fully antiparallel at the metaphase plate. In contrast, the predicted bulk polarity in the assign-to-nearest-pole model (Figure 1D, red curve) predicts that each half-spindle should be composed of almost completely parallel microtubules. This indicates that a simple model in which microtubules branch from the pole is not consistent with bulk spindle polarity as measured by SHG.

### CryoET reconstructions reveal that microtubules in metaphase spindles are arranged in interdigitated, antiparallel bundles with consistent spacing between microtubules

Since our initial efforts to infer microtubule polarity from spindle context in intermediate-resolution serial-section plastic reconstructions were not consistent with bulk polarity fluorescence measurements, we sought to use higher resolution imaging to directly determine the polarity of individual microtubules. Recent advances in cryo-electron microscopy and cryo-electron tomography (cryoEM/cryoET) have revolutionized the field of structural biology and have provided near-atomic-resolution structures of thousands of proteins and complexes. Simultaneous advances in focused ion beam (FIB) technology have enabled the preparation of cellular samples for cryoET by milling away bulk material on either side of a thin lamella^55–57^. Within these lamella, native structures are preserved in vitreous ice at near-atomic resolution.

We prepared metaphase spindles in human RPE-1 cells expressing mCherry-tagged α-tubulin for cryoET. We chose to use RPE-1 cells because they are a human, diploid, non-cancerous cell line^66,67^, which should be representative of human mitotic cells. Briefly, RPE-1 cells were synchronized using a Monastrol^20^ washout (see *Methods*) to enrich cells in mitosis. Prior to milling, we used cryo-fluorescent light microscopy (cryo-FLM) to identify and target cells in metaphase based on the mCherry::α-tubulin label. Areas of interest in metaphase spindles of RPE-1 cells were milled on an Aquilos 2^+^, and tilt series were acquired on the lamella on a Titan Krios (Figure 2A). Tomograms were reconstructed from these tilt series using the WARP^68^-

**Figure 2:**
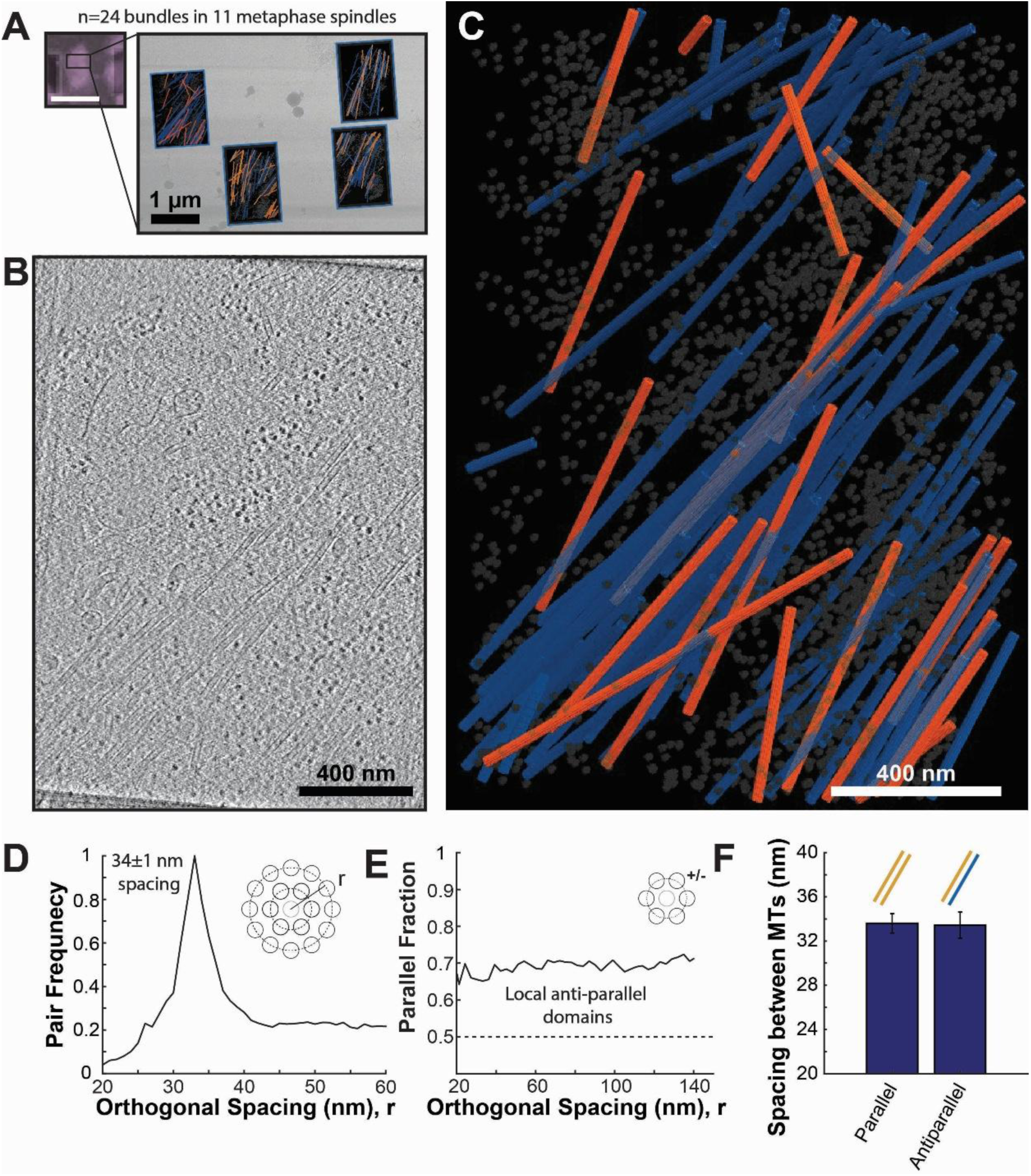
CryoET reconstructions reveal that metaphase microtubules assemble in locally antiparallel domains. A) Cryo-FLM (mCherry::α-tubulin) confirming targeting in a metaphase spindle (left, scale bar 10 µm) and a representative lamella (right) where four tilt series were acquired (total of N=24 tilt series of bundles in 11 metaphase spindles). B) Z-slice of a tomographic reconstruction of a bundle of microtubules in the human metaphase mitotic spindle. Microtubules, ribosomes, and membrane fragments are visible in the tomogram from the lamella in A. C) Annotated segmentation of microtubules and ribosomes in a human metaphase mitotic spindle, shown in B. Orange: microtubules with plus-end pointing toward the top of the image. Blue: microtubules with plus-end pointing toward the bottom of the image. Gray: ribosomes. D) Histogram of spacing between microtubule pairs in bundles in the human metaphase mitotic spindle (orthogonal radial structure factor g(r)). E) Fraction of parallel vs antiparallel microtubules as a function of spacing between microtubules. F) Mean peak spacing between pairs of parallel and antiparallel microtubules (Error bars: SEM).

Aretomo^69^ pipeline (Figure 2B), and microtubules were then initially segmented using the TARDIS-EM segmentation package^70^ followed by manual correction in Amira^71^ (Figure 2C, Figure 2 - supplement 1, Figure 2 - supplement 2).

In these tomograms, we found that microtubules were arranged in dense bundles ∼200-300 nm wide, with low microtubule density in the spaces between the bundles. To investigate the organization of microtubules in these bundles, we computed the orthogonal radial structure factor, g(r), by constructing a histogram of orthogonal distances between microtubules in the bundle^72–75^ (Figure 2D, see *Methods*). We observed a single prominent peak centered at 34±1 nm, indicating a consistent mean spacing between adjacent microtubules in the bundles. This corresponds to a wall-to-wall distance of 8 nm between adjacent microtubules in the bundle lattice^74^.

To further investigate how polar interactions between microtubules and motors assemble these bundles and generate forces that assemble the spindle, we determined the polarity of individual microtubules using subtomogram averaging (see *Methods*). We found that the microtubules in bundles were arranged in locally antiparallel domains with 37±5% of microtubule-microtubule pairs in the bundles antiparallel, roughly consistent with the bulk polarity we observed in second harmonic imaging (Figure 2E). The spacing between parallel and antiparallel microtubules was the same (Figure 2F, Parallel: 34±1 nm; Antiparallel: 33±1 nm; p=0.92 Student’s t-test), suggesting that the spacing between, and consequently the arrangement of microtubules was set by an apolar mechanism rather than directly via crosslinking by different-sized motors with different parallel/antiparallel binding preferences. Taken together, these cryoET reconstructions indicate that microtubules in metaphase spindles are organized in interdigitated, locally antiparallel bundles with consistent spacing between neighboring microtubules.

### Spindle bipolarity is not required for antiparallel bundle formation

Intrigued by these results, we wanted to understand what determines the spacing between microtubules and how this spacing allows binding of motors and crosslinkers of different sizes. We also wanted to further investigate how antiparallel microtubule overlap forms and what determines the local polarity organization. One possibility is that a significant fraction of microtubules from the opposing spindle pole cross over and interdigitate with the locally-generated microtubules to form antiparallel bundles. Alternatively, microtubules could be bundled locally by a motor protein. Lastly, microtubules could be transported from the other half of the spindle; however, we assume that transport is unlikely as the microtubules move too slowly (2.5 µm/min maximum) and turnover is too fast (8 s lifetime), so the average microtubule would only move 300 nm before it disappears^43^. Thus, we assume that the region in the tomographic data where we observe a microtubule is most likely the place where it was nucleated.

To investigate these questions, we sought to determine how the spacing and polarity of microtubules in the spindle responded to molecular motor perturbations. Motor perturbations, however, are typically coupled to changes in spindle geometry^6,19,20^. For example, inhibiting the plus-end-directed Eg5 motor collapses the two opposite spindle poles to form an aster^20,21^, removing both the ability of the motor to locally generate antiparallel interactions and a potential source of antiparallel microtubules crossing over the metaphase plate from the (now collapsed) opposite pole. This complicates the interpretation of an Eg5 knockout experiment because it is unclear whether the inhibition of the motor directly prevents local formation of antiparallel interactions or indirectly removes antiparallel microtubules crossing over from the removed opposite spindle poles.

To decouple changes in spindle geometry from motor activity, we generated motor-active, monopolar spindles by treating the cells for 48 hours with a PLK4 inhibitor, Centrinone^31,32^, that inhibited centriole duplication (Figure 3A). After treatment with Centrinone, the cells only contained a single mother centriole, so they often spontaneously formed monopolar spindles without altering molecular motor activity. This enabled us to investigate whether the antiparallel microtubule interactions that we observed in the metaphase bundles were locally organized by motor activity or had originated from antiparallel microtubules nucleating from opposite poles, either by crossing over the metaphase plate or by repeated branching nucleation events that could generate antiparallel microtubules far from the original mother microtubule at the opposite spindle pole.

**Figure 3:**
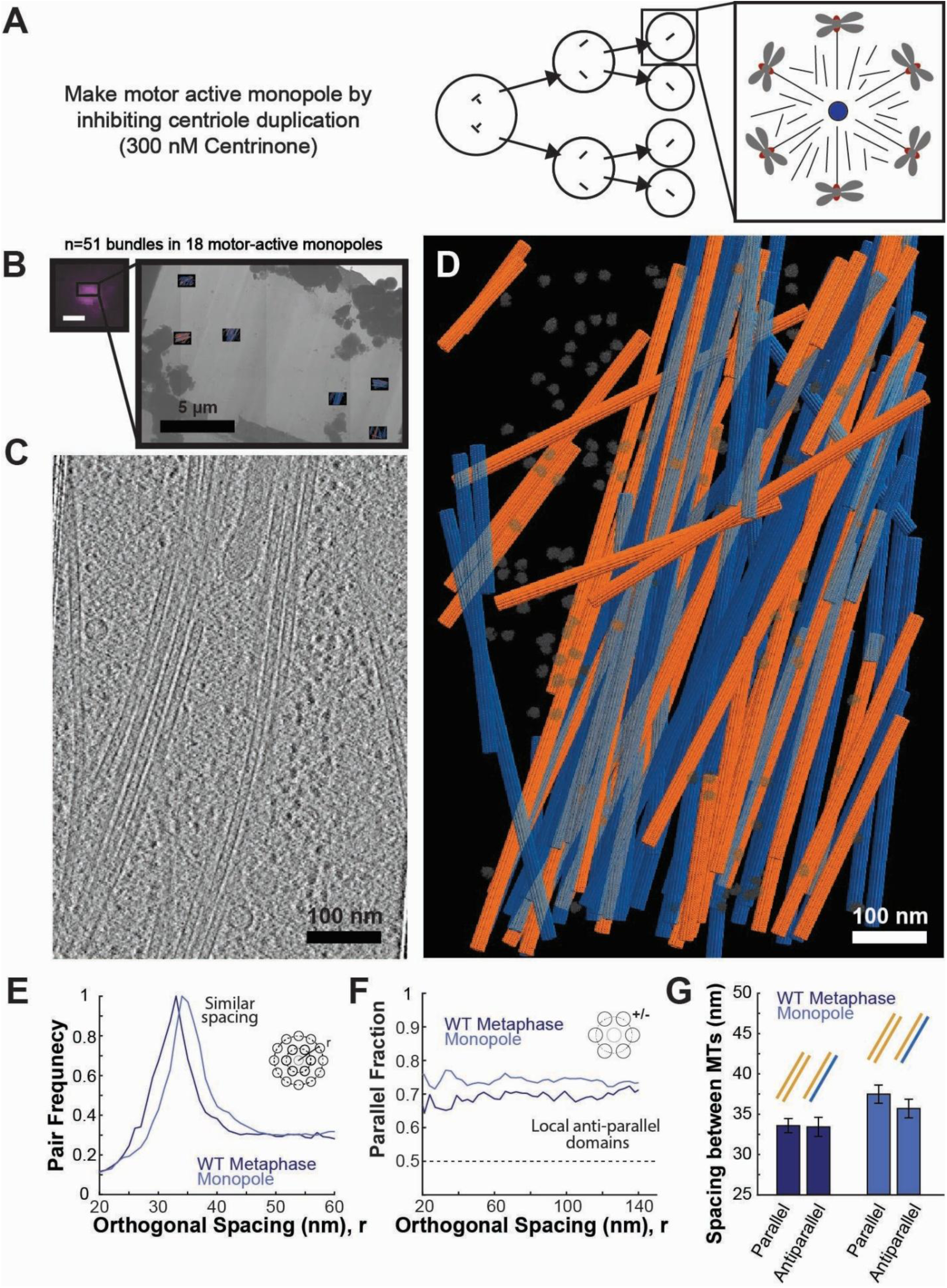
Spindle bipolarity is not required for antiparallel microtubule bundle formation. A) Motor-active monopolar spindles can be generated by inhibiting centriole duplication for two rounds of cell division (300 nM Centrinone for 48 hours in RPE-1 cells). B) Cryo-FLM (mCherry::α-tubulin) confirming targeting of a monopolar spindle (scale bar 2 µm) and a representative lamella, where six tilt series were acquired (Total of N=51 tilt series of bundles in 18 monopolar spindles). C) Z-slice of a tomographic reconstruction of a bundle of microtubules in a motor-active monopolar spindle. D) Annotated segmentation of microtubules and ribosomes of a bundle of microtubules in a motor-active monopolar spindle shown in C. Orange: microtubules with plus-end pointing toward the top of the image. Blue: microtubules with plus-end pointing toward the bottom of the image. Gray: ribosomes. E) Histogram of spacing between microtubule pairs in bundles. F) Fraction of parallel vs antiparallel microtubules as a function of spacing between microtubules. G) Mean peak spacing between pairs of parallel and antiparallel microtubules (Error bars: SEM).

We froze the Centrinone-treated monopolar spindles and confirmed that they were monopolar in cryo-fluorescence (Figure 3B). We then acquired milled, tilt series, reconstructed tomograms (Figure 3C), and segmented and polarity-scored individual microtubules in the tomograms (Figure 3D, Figure 3 - supplement 1). We found that spacing between neighboring microtubules was very similar in the motor-active monopolar and wild-type, bipolar metaphase spindles (Figure 3E, Metaphase: 34±1 nm; Monopole: 36±1 nm; p=0.09 Student’s t-test) and that the bundles still contained a similar fraction of antiparallel microtubule interactions (Figure 3F, Metaphase: 37±5%; Monopole: 28±3%; p=0.09). There was again no statistically resolvable difference between the spacing of parallel and antiparallel microtubule pairs in the motor-active monopoles (Figure 3G; Monopole Parallel: 37±1 nm; Monopole Antiparallel: 35±1 nm; p=0.27 Student’s t-test).

Taken together, these results argue that spindle bipolarity is not required for the formation of antiparallel interactions. Since there is no opposite pole to nucleate antiparallel microtubules in a monopole, the antiparallel microtubules must be locally organized rather than formed by extensive branching nucleation from the opposite pole.

### Inhibition of Eg5 generates parallel bundles in monopolar and bipolar spindles

Since we observed that spindle bipolarity was not required to form antiparallel overlap within bundles, we wanted to test the second hypothesis: that motors can locally organize antiparallel interactions. To do so, we inhibited the two key Eg5 and dynein motors, which reshape the spindle in an aster or a web-like structure, respectively (Figure 4A). Inhibiting both motors restores spindle bipolarity.

**Figure 4:**
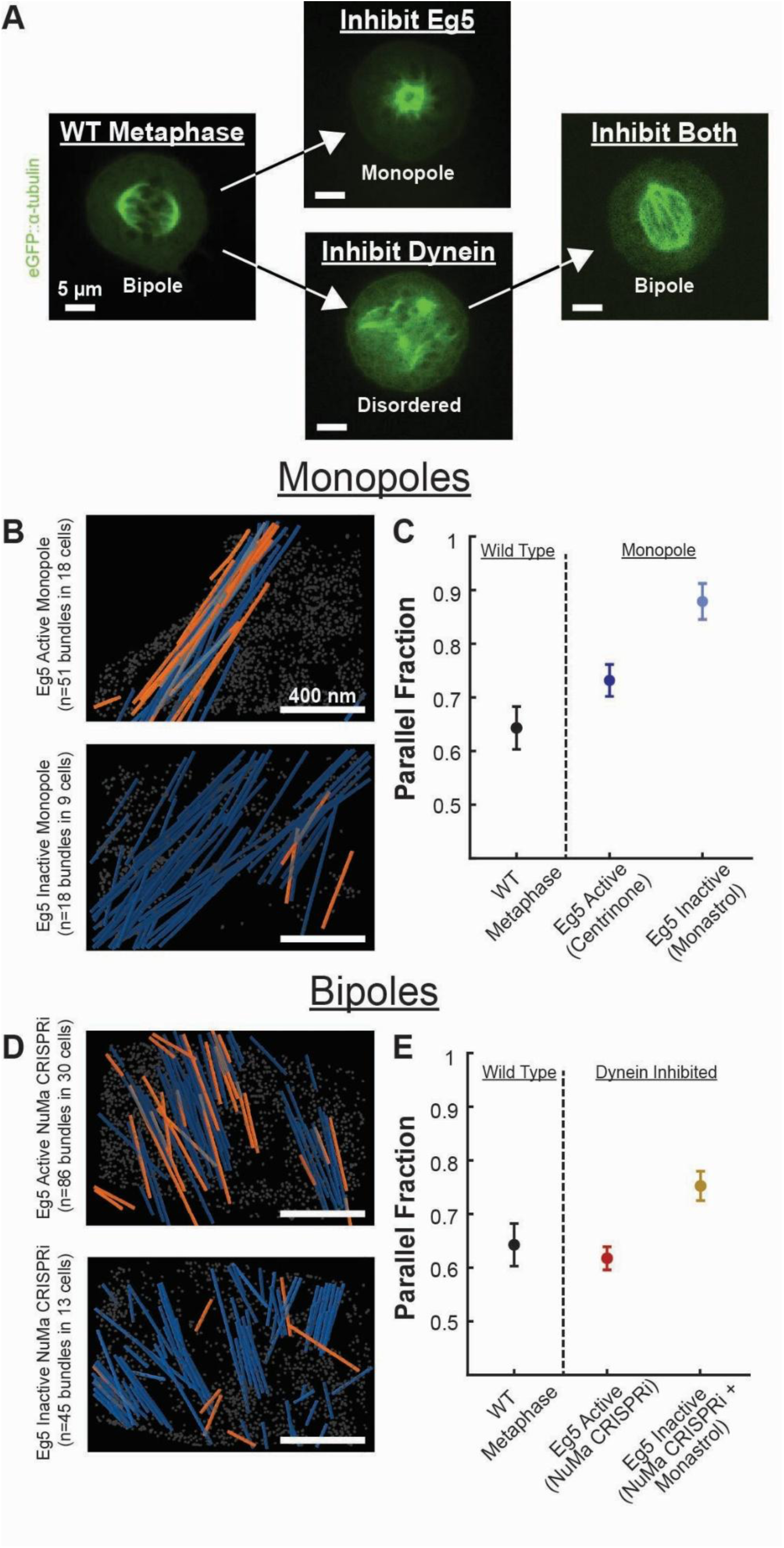
Inhibition of the Eg5 motor generates parallel bundles in monopolar and bipolar spindles. A) Inhibiting Eg5 generates monopolar asters; inhibiting dynein generates disordered, web-like spindles; inhibiting both motors restores spindle bipolarity, B) Representative reconstructions of microtubule bundles in Eg5 active (48 hours in 300 nM Centrinone; N=51 tilt series of bundles in 18 cells) and Eg5 inactive (2 hours in 200 µM Monastrol; N=18 tilt series of bundles in 9 cells) monopolar spindles. Orange: microtubules with plus-end pointing toward the top of the image. Blue: microtubules with plus-end pointing toward the bottom of the image. Gray: ribosomes. C) Mean parallel fraction of nearest neighbors in wild type metaphase, Eg5 active and Eg5 inhibited monopolar spindles (Error bars: SEM). D) Representative reconstruction of microtubule bundles in Eg5 active (N=86 tilt series of bundles in 30 cells) and Eg5 inhibited NuMa CRISPRi dynein inhibited spindles (N=45 tilt series of bundles in 13 cells). E) Mean parallel fraction of nearest neighbors in wild type metaphase, Eg5 active and Eg5 inhibited NuMa CRISPRi dynein inhibited spindles (Error bars: SEM)

We first sought to investigate the effect of Eg5 inhibition. To do so, we treated cells with Monastrol^20,21^, an Eg5 inhibitor, for 2 hours before plunge freezing. After treatment with Monastrol, the bipolar spindles collapsed into a monopole. We again screened spindles with cryo-fluorescent light microscopy (cryo-FLM), milled, acquired tilt series, segmented, and polarity-scored microtubules in these spindles (Figure 4B, Figure 4 - supplement 1A). In contrast to the antiparallel bundles, we observed in the motor-active, Centrinone-treated monopoles, microtubules in the Eg5-inhibited monopoles were significantly more parallel (Figure 4C; Eg5 Active Monopole: 28±3%; Eg5 Inhibited Monopole: 11±3%; p=0.004 Student’s t-test). This indicates that the Eg5 motor locally organized antiparallel bundles in the Eg5 active, Centrinone-treated monopolar spindles.

To further investigate the relationship between spindle bipolarity, antiparallel overlap, and the Eg5 motor, we sought to generate bipolar spindles with Eg5 inhibited. To do so, we first inhibited dynein by depleting NuMa, an adapter protein required to recruit dynein to the spindle, using a previously described NuMa CRISPRi system^19,76^. Then we inhibited Eg5 by treating these spindles with Monastrol to reform an Eg5-inactive bipolar spindle^19^ (Figure 4D). We found that the dynein-inhibited, NuMa CRISPRi spindles had a similar antiparallel fraction to wild-type spindles despite the web-like structure of the dynein-knockout (Figure 4E, WT Metaphase: 37±5%; NuMa CRISPRi: 37±3%; p=0.99 Student’s t-test). In contrast, the dual-inhibited bipolar spindle had more parallel bundles (Dual Inhibited: 25±3%, Meta vs Dual: p=0.04; NuMa CRISPRi vs Dual: p=0.009 Student’s t-test) (Figure 4 - supplement 1B, Figure 4 - supplement 2). Thus, in both monopolar spindles and bipolar spindles, inhibition of the Eg5 motor causes the local arrangement of microtubule bundles to become more parallel, again indicating that Eg5 activity, not spindle bipolarity, is required to form antiparallel microtubule interactions.

### A balance of Eg5 and dynein sets microtubule spacing and organization by regulating microtubule density

Next, we wanted to better understand what determines the spacing between microtubules, and consequently, the organization of microtubule bundles. In wild-type spindles, we found a consistent 8 nm wall-to-wall spacing between microtubules with no difference between parallel and antiparallel pairs. It is not clear what determines this spacing or why there is no dependence on microtubule polarity. One possibility is that microtubules are directly organized via crosslinking by molecular motors or passive crosslinkers, as observed in budding yeast spindles^62^. This would likely be a single, apolar crosslinker, since we found a single apolar peak spacing. In this model, microtubule spacing would be independent of the density of microtubules in the bundle because spacing would instead be set by the length of the crosslinker. Alternatively, bundles could be organized indirectly by density-driven steric interactions between microtubules. This would produce a consistent spacing between microtubules that would scale with density, as observed in nematics^88,89^ or a simple 2D liquid^72,75^.

We first investigated whether the crosslinker model could explain microtubule organization in bundles by examining how the spacing between microtubules changed when the two key motors, Eg5 and dynein, were perturbed (Figure 5A, B). If either of these motors were the organizing crosslinker, then the spacing would change when the motors were perturbed. We found that inhibiting dynein, but not Eg5, increased the spacing between adjacent microtubules (Figure 5B; WT: 34±1 nm, Eg5-inhibited: 32.5±0.4 nm, p=0.2 Student’s t test; NuMa CRISPRi: 40±2 nm, p=0.03 Student’s t test); however, inhibiting both motors restored the wild-type spacing (Dual KO: 35±1 nm, p=0.6 Student’s t test). This excludes the possibility that another crosslinker besides dynein determines the microtubule spacing because wild-type spindles had a single peak spacing, consistent with a single crosslinker organizing the bundles, and inhibiting dynein altered spacing. If microtubule bundles were organized by direct crosslinking by dynein, however, then the spacing would not return to wild-type levels when both motors were inhibited. This indicates that dynein did not directly crosslink microtubules in the bundles and that the crosslinker model was not consistent with changes in microtubule spacing after motor inhibitions.

**Figure 5:**
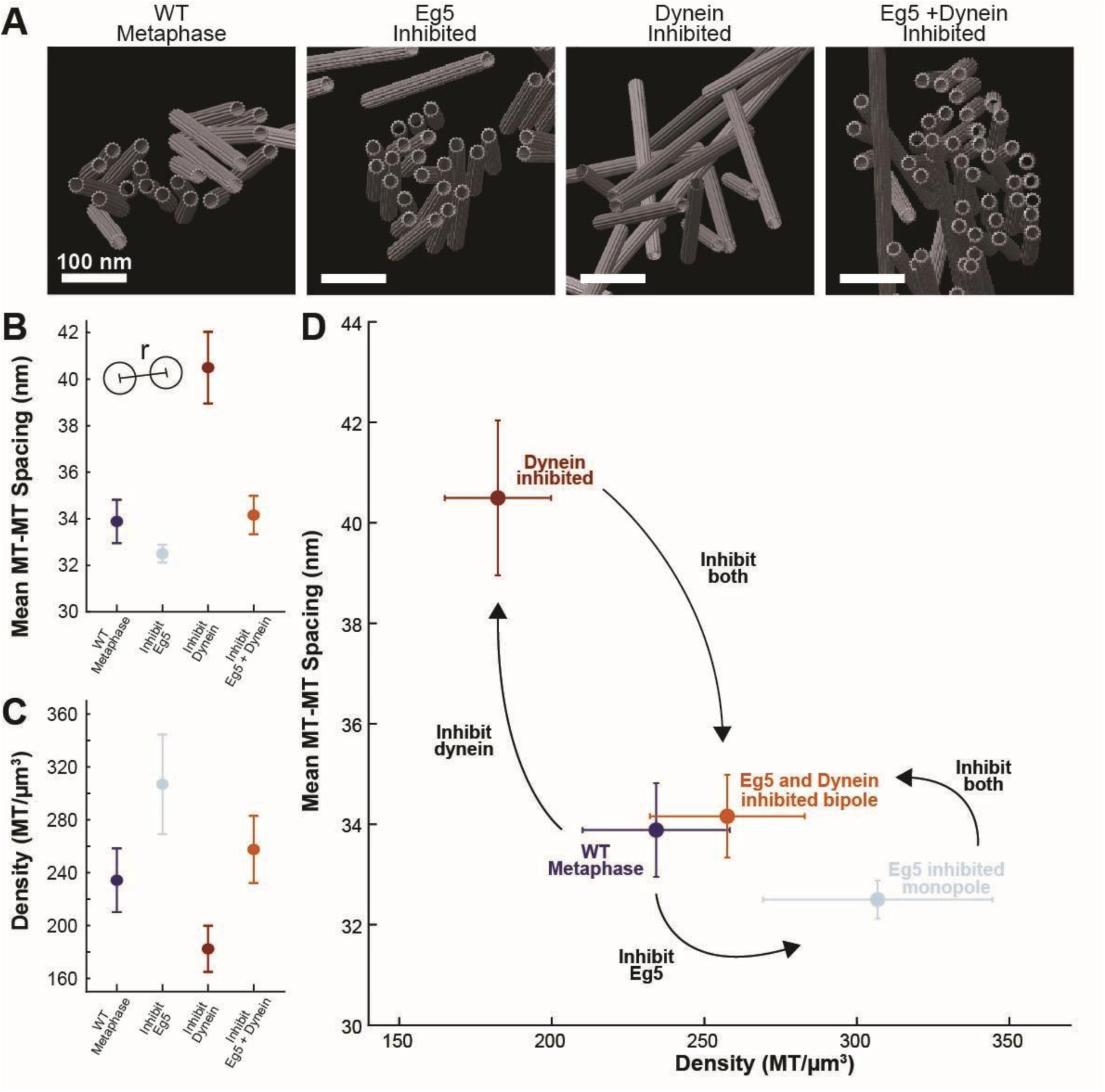
A balance of Eg5 and dynein regulates microtubule spacing and density. A) Side-views of cryoET reconstructions of microtubule bundles in wild-type metaphase, Eg5-inhibited monopolar spindles, dynein-inhibited NuMa CRISPRi-treated spindles, and dual-inhibited spindles. B) Mean peak spacing between microtubule centerlines in each bundle condition. C) Mean microtubule density (microtubule/µm³) in each bundle condition. D) Mean microtubule-microtubule peak spacing vs density for each motor knockout condition.

We then investigated whether the second model, that density-driven steric interactions organize microtubule bundles, was consistent with microtubule spacing in the motor-perturbed spindles. To do so, we calculated the mean density of microtubules in each of the motor conditions (Figure 5C) and then plotted how the spacing varied with density for each condition (Figure 5D). We found that inhibiting dynein decreased microtubule density and significantly increased microtubule spacing, whereas inhibiting Eg5 slightly increased microtubule density and decreased spacing. Inhibiting both motors restored WT spacing and density, suggesting the motor inhibitions regulated microtubule spacing by altering microtubule density rather than by removing possible crosslinkers that directly set microtubule spacing.

To gain further insight into the relationship between microtubule spacing and local density, we returned to the lower-resolution, full-spindle plastic reconstructions of wild-type HeLa metaphase spindles (Figure 6A). These reconstructions captured a much larger total volume (640 µm^3^ per spindle vs ∼0.3 µm^3^ per cryotomogram), providing increased data coverage, particularly in lower density regions between bundles which we did not target when collecting cryotomograms of microtubule bundles in spindles. We found that the spacing between microtubules was similar in the cryo and plastic reconstructions (Figure 6 - supplement 1A). To account for expansion/shrinkage in the plastic, we normalized the three plastic spindle reconstructions to have the same peak spacing as the cryotomograms so that they could be compared against one another (see *Methods*). This normalization process does not alter the scaling between spacing and density. We found that the spacing between adjacent microtubules was higher in the center of the spindle, where the density is lower, than near the spindle poles (Figure 6B). We collapsed the data from all regions into a single spacing vs. density plot and found that the spacing between microtubules decreased as density increased in all three plastic spindles independently (Figure 6 - supplement 1B) and when the spindles were normalized and averaged together (Figure 6C). This relationship between density and spacing is not consistent with a purely crosslinker model where spacing is independent of density and set by the length of the crosslinker and is consistent with a model where spacing is set purely by density-driven, steric interactions between microtubules (as predicted by a simple liquid crystal model^72,75,77,78^).

**Figure 6:**
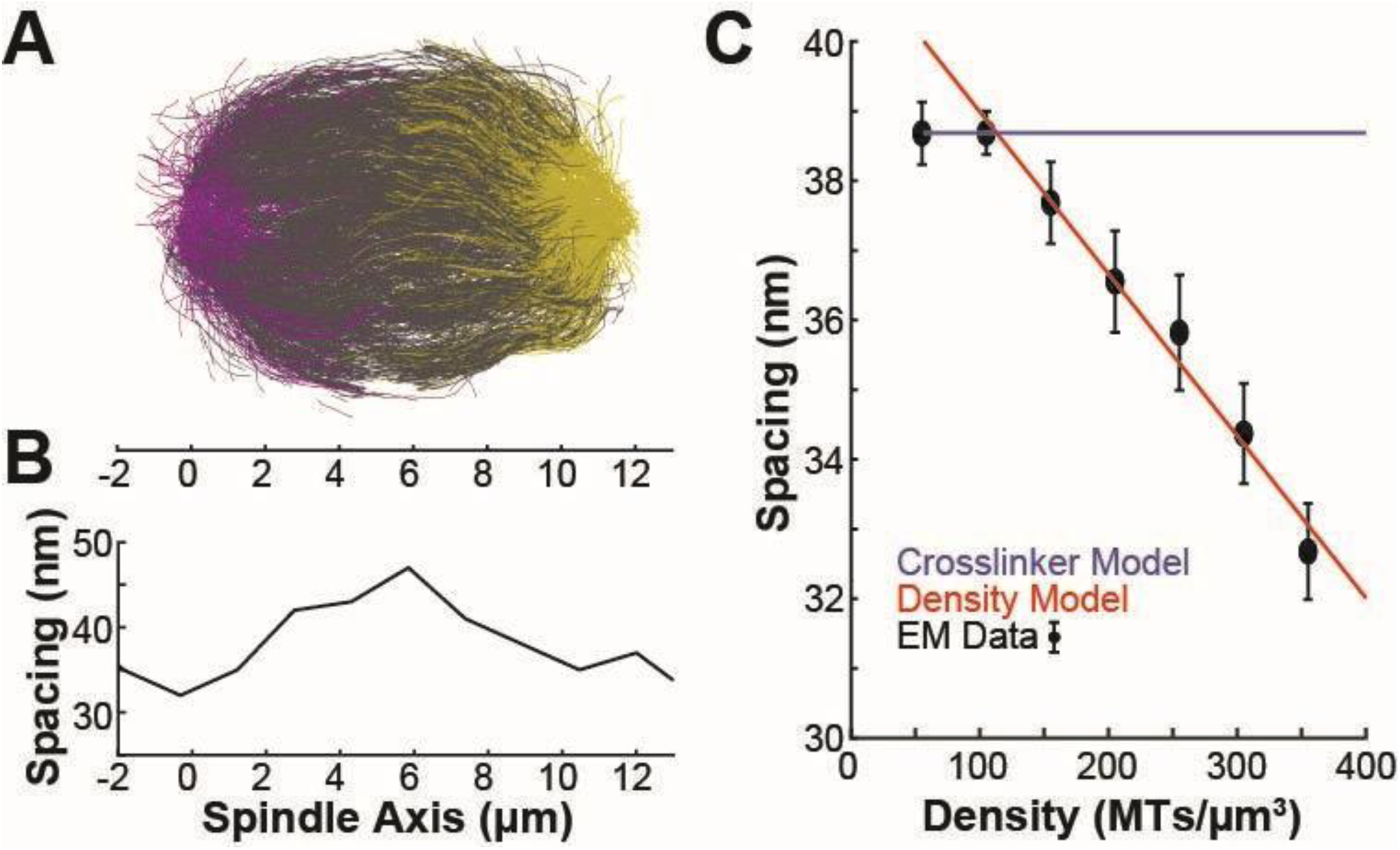
Microtubule density sets spacing between microtubules in the human metaphase mitotic spindle. A) Reconstruction of the trajectory of every microtubule in a metaphase HeLa spindle (Kiewisz et al. 2022). B) Mean peak spacing between microtubules along the spindle axis of the spindle in A. C) Spacing between microtubules (normalized to account for differences in contraction in the resin in different reconstructions) vs microtubule density. Blue: Prediction from a model where spacing is set by a single crosslinker. Red: Prediction from a model where spacing is set by microtubule density. Black: Data from reconstructions.

## DISCUSSION

Here, we utilized recent advances in cellular cryo-electron tomography^55–57^ to investigate how polarity-dependent interactions between molecular motors and microtubules assemble the mitotic spindle. Previous reconstructions of the spindle have utilized resin-embedded electron tomography, which compromises imaging resolution, so the polarity of individual microtubules cannot reliably be determined^41,42,53,54^. This makes it difficult to investigate how polar interactions between microtubules and motors assemble the spindle. To overcome this, we initially attempted to assign microtubule polarity in these low-resolution reconstructions based on the microtubules’ locations in the spindle, assigning each microtubule to the nearest pole (Figure 1). We found that this simple model was not consistent with SHG measurements of bulk microtubule polarity in the spindle, so we turned to higher-resolution cryoET imaging to resolve the microtubule polarity directly from electron microscopy. From this imaging, we found that microtubules organize in dense bundles with local antiparallel overlap and consistent 8 nm wall-to-wall spacing between adjacent microtubules (Figure 2). Using a combination of centriole duplication and motor inhibitions, we determined that the antiparallel microtubule overlap was generated by the Eg5 motor, not by microtubules branching from the opposite spindle pole (Figures 3 and 4). We found no significant difference in spacing between parallel and antiparallel microtubule pairs, implying that spacing is set by an apolar process rather than by direct crosslinking by polarity-dependent molecular motors. Instead, spacing scaled with microtubule density in cryotomograms taken in both wild-type and motor-mutant spindles (Figure 5) and in large-scale plastic reconstructions (Figure 6).

These results point to a simple, multiscale mechanistic picture of spindle assembly driven by a balance of the Eg5 and dynein motors at the level of individual motors and microtubules, microtubule bundles, and the entire spindle. At the level of individual motors and microtubules, Eg5 crosslinks and slides antiparallel microtubules, while dynein clusters microtubule minus-ends as cargo (Figure 7A). At the bundle scale, Eg5 arranges antiparallel overlap, which generates outward extensile forces that separate the spindle poles. Dynein balances these extensile forces to increase microtubule density and maintain consistent spacing within the bundle. At the scale of the entire spindle, Eg5 provides extensible active stress, which balances surface tension generated by dynein to form an active, ellipsoidal, bipolar structure.

**Figure 7:**
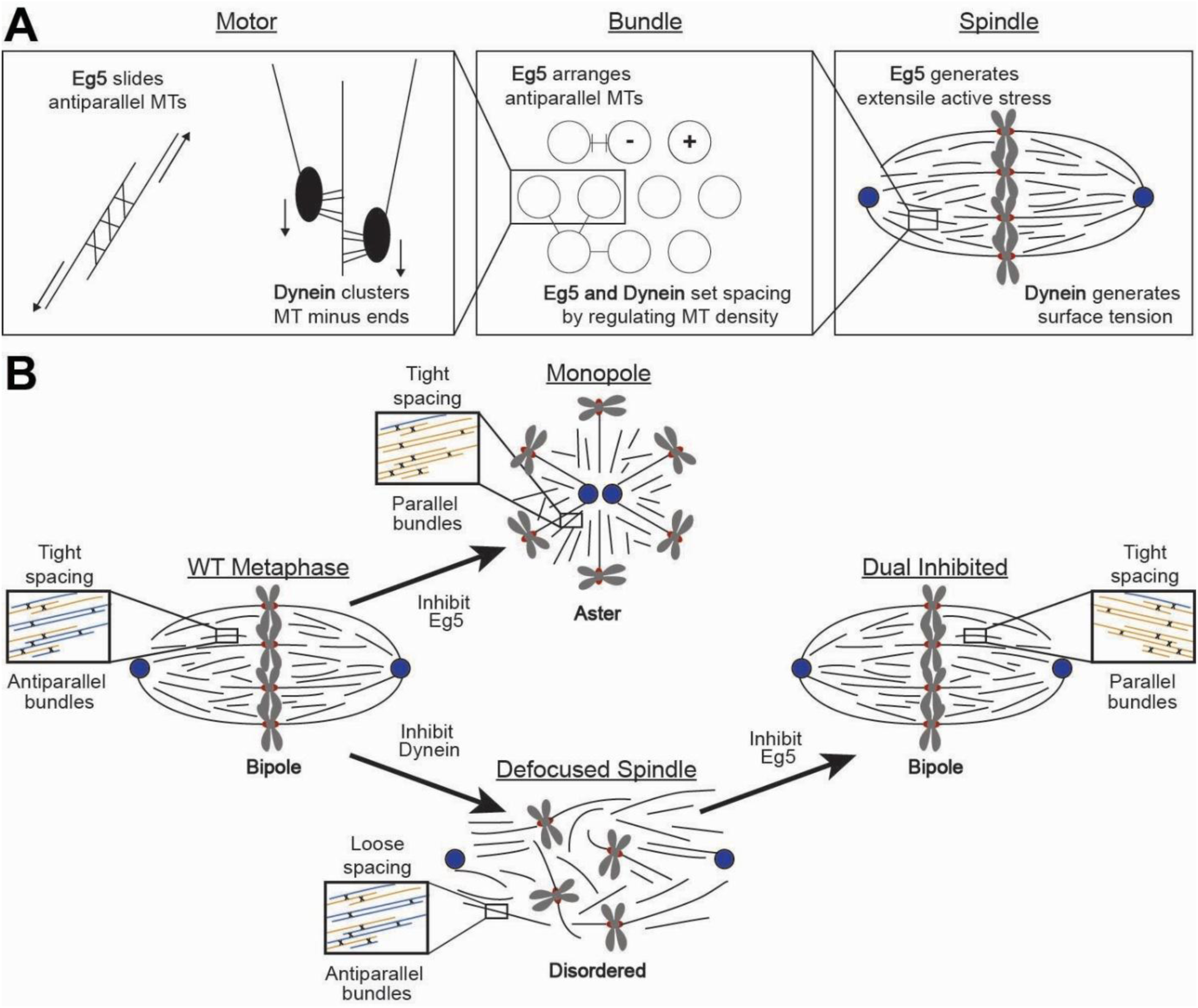
A multiscale picture of spindle assembly. A) At the motor scale, Eg5 slides antiparallel microtubules while dynein clusters microtubule minus-ends by transporting microtubules to minus-ends as cargo. At the bundle scale, Eg5 arranges antiparallel microtubule pairs while a balance of Eg5 and dynein sets spacing by regulating microtubule density. At the spindle scale, Eg5 generates extensile active stress to separate the spindle poles while dynein generates a contractile surface tension. B) In bipolar, WT metaphase spindles, bundles are organized with antiparallel overlap and tight spacing. Inhibiting Eg5 generates a monopolar aster with parallel bundles and tight spacing. Inhibiting dynein generates a defocused spindle with antiparallel bundles and loose spacing. Dual inhibition of Eg5 and dynein refocuses the spindle poles to form a bipolar spindle with tight, parallel bundles.

Inhibiting Eg5 generates parallel bundles that cannot generate extensile outwards stresses, so the spindle poles collapse to form an aster (Figure 7B). Inhibiting dynein reduces microtubule density, so bundles no longer form, defocusing the spindle poles into a web-like structure composed of antiparallel bundles assembled by Eg5. Inhibiting both motors increases the spindle density, reforming tight bundles. Without Eg5, however, the bundles are composed of more parallel microtubules. These spindles can still enter anaphase, though they are much less resistant to mechanical stress and segregate chromosomes less robustly^19^.

In this view, microtubule organization is not determined by the overall spindle shape, with microtubule nucleation exclusively at the poles. Instead, motor proteins act bottom-up by locally organizing microtubules and the spindle poles set up an architecture that enables the motors to interact with microtubules. We propose that the kinesin-5 motor, Eg5, locally creates antiparallel microtubule bundles independently of broader spindle structure. This is supported by three independent lines of experimental evidence: 1) the bulk polarity of microtubules, as measured by both SHG and cryoET, cannot be explained by a branching-from-pole model alone, and microtubules move too slowly to be transported from one spindle half to the other during their lifetime; 2) antiparallel, interdigitated bundles formed in monopolar spindles with active Eg5 (Centrinone); and 3) inhibition of Eg5 reduced the formation of antiparallel bundles within monopolar (Monastrol) and bipolar (dualKO) spindles, indicating that spindle geometry does not affect microtubule polarity. To organize these antiparallel bundles, Eg5 may grab short stubs of freshly nucleated microtubules, either nucleated by Augmin in the spindle bulk^36–40^ or by Ran near the chromosomes^79–82^, and align them in an antiparallel orientation. Alternatively, Eg5 itself could contribute to microtubule nucleation^83^. Further experiments or *in vitro* experiments with purified components would be very informative in better understanding how Eg5 activity organizes antiparallel overlap in spindles.

We found that spindle microtubules organized in dense bundles with microtubules ∼5x more likely to be found at 8 nm wall-to-wall spacing than at any other distance. We also found no difference in spacing between parallel and antiparallel microtubules. We found that this spacing was set by apolar density-driven steric interactions between microtubules rather than directly by motor crosslinking. We observed four direct lines of experimental evidence supporting this claim: 1) there is only a single peak in the orthogonal structure factor rather than multiple peaks corresponding to multiple motors; 2) there was no difference in spacing between parallel and antiparallel microtubules; 3) though inhibiting dynein increased spacing, inhibiting both Eg5 and dynein restored wild-type spacing, indicating that dynein was not responsible for directly crosslinking microtubules; 4) spacing scaled inversely with density across all conditions in cryotomograms and in large scale plastic reconstructions. In this view, the Eg5 and dynein motors act interchangeably in regulating microtubule density, though only the tetrameric, antiparallel Eg5 can regulate microtubule polarity. These bundles could be held together by depletion forces from the viscous cytoplasm, though further theoretical work and experiments in cells with perturbed viscosities will be required to investigate the origin of these density-dependent bundling forces.

It is not immediately clear how motors navigate these tight, bundle lattices. The spacing between microtubules is very narrow (∼8 nm wall-to-wall between microtubules) and most motors are too large to fit in such a narrow gap. The motors could possibly localize in gap defects, where microtubules are missing in the lattice structures, so the motors can have more room to maneuver. Alternatively, the motors could be localized in a ring-like shell on the edge of the bundles. These bundles could resemble a coaxial cable with an older, bundle core inaccessible to the motors. On the bundle edge, freshly recruited microtubules could freely interact with motors and flip around to form antiparallel pairs. Finally, the motors could crosslink non-adjacent microtubules in the second or even third coordination shell, where there would be more spacing between the microtubules. To understand interactions between microtubules and motors in these dense lattices, it will be crucial to develop methodology to directly visualize motors interacting with microtubules in dense bundles both *in vitro,* where the action of individual motors can be directly tested, and in cells, where the native structure can be visualized^84,85^. For smaller or narrow microtubule binders, such as Eg5 or PRC1, larger tags akin to a GFP tag in fluorescent microscopy would be very informative, though for larger motors such as dynein visualization should be possible directly from tomography without tagging.

In the future, it will be essential to combine different approaches that provide information about live cell dynamics, microtubule architecture, and the molecular properties of microtubules to bridge structural and cell biology. While cryoET has the potential to deliver molecular resolution of microtubules in the spindle and may be able to help visualize molecular motors or microtubule crosslinkers *in situ*, it is currently limited to small, thin sections of ∼150 nm thickness within a cell. This physical restriction of the field of view eliminates the spindle context, making it difficult to determine where in the spindle the tomogram is located and where the microtubules originate. Using serial sectioning of resin-embedded spindles, complete reconstructions can be generated at ∼2 nm resolution, providing contextual information at the loss of higher resolution that would allow, for example, the quantification of polarity. While each of these approaches alone is powerful, the combination and the ability to obtain data from nano- to micrometer scale will be key to developing a clear and correct understanding of spindle assembly and function.

## DATA AVAILABILITY

Raw tomograms will be available on EMPIAR at time of final publication. Fluorescence microscopy images will be available on Dryad at time of final publication. Cell lines are available on request from the corresponding author.

## ACKNOWLEDGMENTS

We thank the NCITU milling team, particularly Mykhailo Kopylov, for technical assistance in preparing mitotic cells for cryoET and time on the Aquilos needed to mill samples. We thank Gloria Ha, Dan Needleman, Lila Neahring, and Sophie Dumont for generously providing the cell lines we used in this study. Jake Johnston and Reza Paraan’s advice on cryoET data processing was instrumental to the reconstruction and analysis of the tomograms. Robert Kiewisz’s development of the TARDIS-EM segmentation package single-handedly cured Will’s carpal tunnel syndrome, for which he is immensely grateful.

We have been fortunate to have many productive discussions with collaborators over the course of this project, and we, in particular, would like to thank Dan Needleman, Thomas Mueller-Reichert, Mike Shelley, Reza Farhadifar, Bryce Palmer, Chris Edelmaier, Colin McManus, and Yong Song for their sharp insights and advice.

A Simons Foundation CCB_x_ 00012463 grant supported Will Conway for the duration of this project. Stefanie Redemann was supported by NIH NIGMS 1R01GM144668, Stefanie Redemann and Vitaly Zimyanin were supported by Simons Foundation CCBx 01157393. Alex de Marco was supported by NIH NIGMS 1R24GM154192 and Simons Foundation International grant 586810. Daija Bobe was supported by NIH U24GM139171.

Some of this work was performed at the National Center for In-Situ Tomographic Ultramicroscopy (NCITU) and the Simons Electron Microscopy Center located at the New York Structural Biology Center, supported by the NIH Common Fund Transformative High Resolution Cryo-Electron Microscopy program (U24 GM129539,) and by grants from the Simons Foundation (SF349247) and NY State Assembly

## METHODS AND MATERIALS

### Key resources table

| REAGENT or RESOURCE | SOURCE | IDENTIFIER |
| --- | --- | --- |
| Chemicals, peptides, and recombinant proteins |  |  |
| Monastrol | Selleckchem | - |
| Centrinone | Selleckchem | - |
| Doxycycline | Selleckchem | - |
| Deposited data |  |  |
| Plastic reconstructions of human metaphase spindles | Mueller-Reichert Lab, TU Dresden | - |
| Experimental models: Cell lines |  |  |
| Wild-type HeLa Kyoto cells | Needleman Lab, Harvard | CVCL_1922 |
| HeLa Kyoto + GFP:: α-tubulin | Needleman Lab, Harvard | CVCL_1922 |
| RPE-1 cells + sfGFP::CENP-A, | Needleman Lab, Harvard | CVCL_4388 |
| RPE-1 cells + dox-inducible NuMa CRISPRi + eGFP :: α-tubulin | Dumont Lab, UCSF | CVCL_4388 |
| Software and algorithms |  |  |
| WARP 1.10beta | Genentech | - |
| Aretomo 1.3.0 | UCSF/CZII | - |
| Relion 4 | Scheres Lab | - |
| Matlab 2022b | Mathworks | - |
| TARDIS-em v0.3.10 | SMLC | - |

### Cell culture and sample freezing

RPE-1 cells were thawed from aliquots and cultured in DMEM (ThermoFisher) supplemented with 10% tetracycline-free FBS (ThermoFisher) and Pen-Strep (ThermoFisher) in a humidified incubator at 37 °C with 5% CO_2_. Cells were regularly tested for mycoplasma contamination (Southern Biotech). The RPE-1 cell line expressing mCherry::α-tubulin was a gift from Gloria Ha and Daniel J. Needleman^66^. The RPE-1 cell line expressing GFP::α-tubulin and the dox-inducible NuMa CRISPRi system was a gift from Sophie Dumont^19^. RPE-1 cells are female.

To generate wild-type metaphase spindles, cells expressing mCherry::α-tubulin were initially plated in T175 cell culture flasks to 70% confluency. Cells were synchronized in mitosis via an initial shakeoff to remove dead cells, followed by a two-hour incubation with 200 µM Monastrol (Seleckchem) to arrest cells in mitosis. After this incubation, mitotic cells were detached from flasks via a second mitotic shakeoff, concentrated via centrifugation at 400*xg*, resuspended, and then plated onto poly-l-lysine (ThermoFisher) coated Au 200 mesh holey gold UltrAuFoil EM grids (Quantifoil). Samples were incubated for 5 minutes, then washed into fresh, untreated culture media (0 µM Monastrol) and subsequently incubated for 20-30 further minutes to reform bipolar spindles. Samples were then frozen either by plunge-freezing into liquid ethane on a Leica GP2 plunge freezer or by high-pressure freezing on a Wohlwend Compact 01 using the Waffle Method^86^. Just prior to freezing, cells were incubated in fresh media containing a 5% glycerol cryoprotectant for 30 seconds.

Motor mutant and centriole-depleted spindles were frozen in the same manner with the following changes:

- To generate motor-active, monopolar spindles, cells were treated with 300 nM Centrinone (Seleckchem) for 48 hours prior to the first mitotic shakeoff and subsequently incubated in fresh media without Monastrol for all steps until frozen.
- To generate Eg5-inhibited, monopolar spindles, cells were incubated in media containing 200 µM Monastrol after plating on EM grids (i.e., the Monastrol was never washed out).
- To generate dynein-inhibited, NuMa CRISPRi spindles, cells were incubated in 2 µM doxycycline for 4 days prior to the initial mitotic shakeoff and were incubated in fresh media without Monastrol for the entire shakeoff-incubation-shakeoff procedure until frozen.
- To generate dual-inhibited, bipolar spindles, the dynein-inhibited cells were incubated in media containing 200 µM Monastrol after plating on EM grids for 20-30 minutes.

### FIB-milling and tilt series acquisition

Cells were screened on a Zeiss Axio Imager 2 microscope to identify cells in the correct stage of mitosis (metaphase bipole, monopole, disordered) in the tubulin channel (eGFP ex: 488 nm em: 484-630 nm; mCherry ex: 561 nm em: 567-700 nm) with a 10x long working distance air objective (NA=0.4). Cells on grids were maintained at -194 °C to prevent devitrification on a Linkam CMS196v³ cryostage during imaging.

After fluorescent screening, cells were milled on an Aquilos 2^+^ FIB-SEM microscope (ThermoFisher/FEI). Cryo fluorescence images were overlaid on an initial scanning electron microscope montage image of the grid to locate mitotic cells. Grids were then coated with GIS and automatically milled using AutoTEM (ThermoFisher/FEI) with the following parameters: Initial rough mill: 1 nA, Medium mill: 1 nA; Fine milling: 0.5 nA; Finer milling: 0.3 nA; Polishing 1: 100 pA, Polishing 2: 50 pA. After automated milling, lamellae were manually cleaned to remove defects left by automation. Grids frozen using the Waffle Method were milled as described previously. AutoTEM milling templates are available upon request from the corresponding author.

Milled grids were then loaded into a Titan Krios (FEI/Thermo Fisher) equipped with a K3 direct electron detector camera (Gatan) and a BioQuantum Imaging Filter at NCITU NYSBC for tilt series collection. Tilt series were collected using SerialEM with 3 ° dose-symmetric increments at 3 µm target defocus and a dose of 3.1 e/Å^2^ per tilt from ±50 ° from the milling angle (if the milling angle was higher than 10 °, as in the Waffle Method, then the tilt range was truncated early at 60 °). Tilt series were collected at both 26,000x (3.4 Å/pixel) in 0.5x superresolution mode and 42,000x in standard 1x counting mode (2.07 Å/pixel). SerialEM settings files for acquisition are available upon request from the corresponding author.

### Tomogram reconstruction, segmentation, and microtubule polarity determination

Following data acquisition, individual tilts were preprocessed for CTF estimation and motion correction in WARP 1.10beta^68^ before export to Aretomo1.7^69^ for tilt series alignment. Tilt series were manually pruned to remove poorly aligned tilts. After alignment, tomograms were reconstructed in WARP, and microtubules were segmented using the TARDIS-EM segmentation package^70^ followed by manual correction in Amira^71^. Subtomograms were then extracted every 4 nm down the microtubule trajectories in WARP.

To determine the microtubule polarity, we performed subtomogram averaging in Relion4 ^87^ (Figure 2 - supplement 1). We generated an initial microtubule model *ab initio* and then performed four total rounds of refinement: C1 and then helical with 10 Å pixels, followed by C1 and then helical with 4 Å pixels. Each motor mutant condition was processed separately. Following subtomogram averaging, polarity was assigned by consensus along the microtubule^53,88^ (i.e., assigning the polarity as the direction that the majority of the subtomograms pointed).

### Second harmonic imaging

We utilized a HeLa Kyoto cell line expressing GFP::α-tubulin which was a kind gift from Daniel J. Needleman. HeLa Kyoto cells are female. Cells were prepared for imaging by plating on poly-d-lysine-coated coverslips one day prior to imaging. 10 minutes prior to imaging, cells were incubated in imaging media containing Fluorobrite DMEM (ThermoFisher) supplemented with 10 mM HEPES and then transferred to a custom heater where they were maintained at 37 °C during imaging. Second harmonic images were acquired on a custom-built two-photon microscope with an Insight X3 Ti-Sapphire 80 MHz pulsed laser (Spectra Physics) at 850 nm. The incoming photons were circularly polarized by a half- and quarter-wave plate setup to ensure spatially uniform SHG detection. Second harmonic and GFP:: α-tubulin images were acquired simultaneously in the transmission and reflection arms, respectively (SHG: 425nm/30 nm + 650 nm short-pass; GFP: 535 nm/30 nm + 650 nm short-pass).

Bulk spindle polarity was measured from the second harmonic images by first projecting the SHG and tubulin intensities onto the spindle axis. These intensities were corrected for background by subtracting off the mean SHG/GFP::tubulin signal outside of the spindle in the cytoplasm. Bulk polarity along the spindle axis was calculated from these curves via the relationship 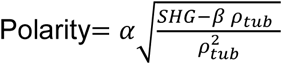, where 𝜌_𝑡𝑢𝑏_ is the density of tubulin, SHG is the second harmonic intensity, and the 𝛼 and 𝛽 constants are set by the spindle boundary conditions as previously described and validated^65^.

### Whole-spindle serial-section resin-embedded electron tomography

We utilized a previously published reconstruction of three complete metaphase spindles in HeLa cells (Kiewisz et al. 2022) available [here](https://opara.zih.tu-dresden.de/xmlui/handle/123456789/5675).

### Polar bundle lattice structure analysis

To analyze the lattice structure of the microtubule bundles, we first spline fit the trajectories of the microtubules using the microtubule center points obtained after subtomogram averaging. For each of these microtubule trajectories, we defined a plane orthogonal to the microtubule trajectories every 1nm along the trajectory. We then drew a vector in this orthogonal plane from the microtubule trajectory centerline to the center of all other microtubules that crossed the plane. We defined the orthogonal distances between the microtubules as the magnitudes of these orthogonal plane vectors between microtubules and computed a histogram of these distances to calculate the radial distribution function, g(r) (Figure 2D, 3E). To calculate the angles between neighboring microtubules in the bundles, we computed the angles between these orthogonal plane vectors for microtubules in the first coordination shell of the bundle, defined as 25-45nm orthogonal spacing from the original microtubule (Figure 5C).

We assigned the polarity of microtubules in the bundle by first determining the overall orientation of the bundle by calculating the mean nematic director field of the microtubules by calculating the mean nematic stress tensor, Q, for all microtubules in the bundle^77,78^. We then projected the polar microtubule trajectory vectors, determined by mean consensus along the microtubule after subtomogram averaging, onto the mean nematic director field and assigned the polarity of the microtubules as “forwards” or “backwards” relative to one another. To calculate the mean parallel fraction of microtubules in the bundle, we calculated how many pairs of microtubules in the first coordination shell, again defined as 25-45 nm orthogonal spacing, pointed in the same (parallel) or opposite (antiparallel) directions.

To analyze how spacing varied with microtubule density in the plastic reconstructions, we subdivided the complete spindle reconstructions into smaller 1 µm x 1 µm x 0.2 µm slices and ran the same spacing analysis as we did for the cryo-tomograms. We initially binned each of these regions together for each spindle to calculate the peak spacing for each spindle independently (Figure 6 – supplement 1A) and found that the peak spacing was very similar to the cryotomograms for two of the plastic spindles, but the third spindle peak spacing was slightly higher. We presume that this difference in spacing occurred because of variable expansion/shrinkage during resin-embedding and acquisition, so we normalized all three spindles to have the same peak spacing as the untreated cryotomograms to compare how spacing scaled with microtubule density in the three spindles (Figure 6C). We found, however, that the spacing decreased with increased microtubule density in all three plastic spindles when analyzed independently (Figure 6 – supplement 1B).

### Quantification and Statistical Analysis

In all figures and in the main text, error bars and ± symbols are standard errors of the mean. The number of bundles included in each condition is included in the figures where the experiment is introduced and in the figure captions. Spacing and polarity for all conditions were compared using the Student’s t-test.

---

**Figure 1 - supplement 1:**
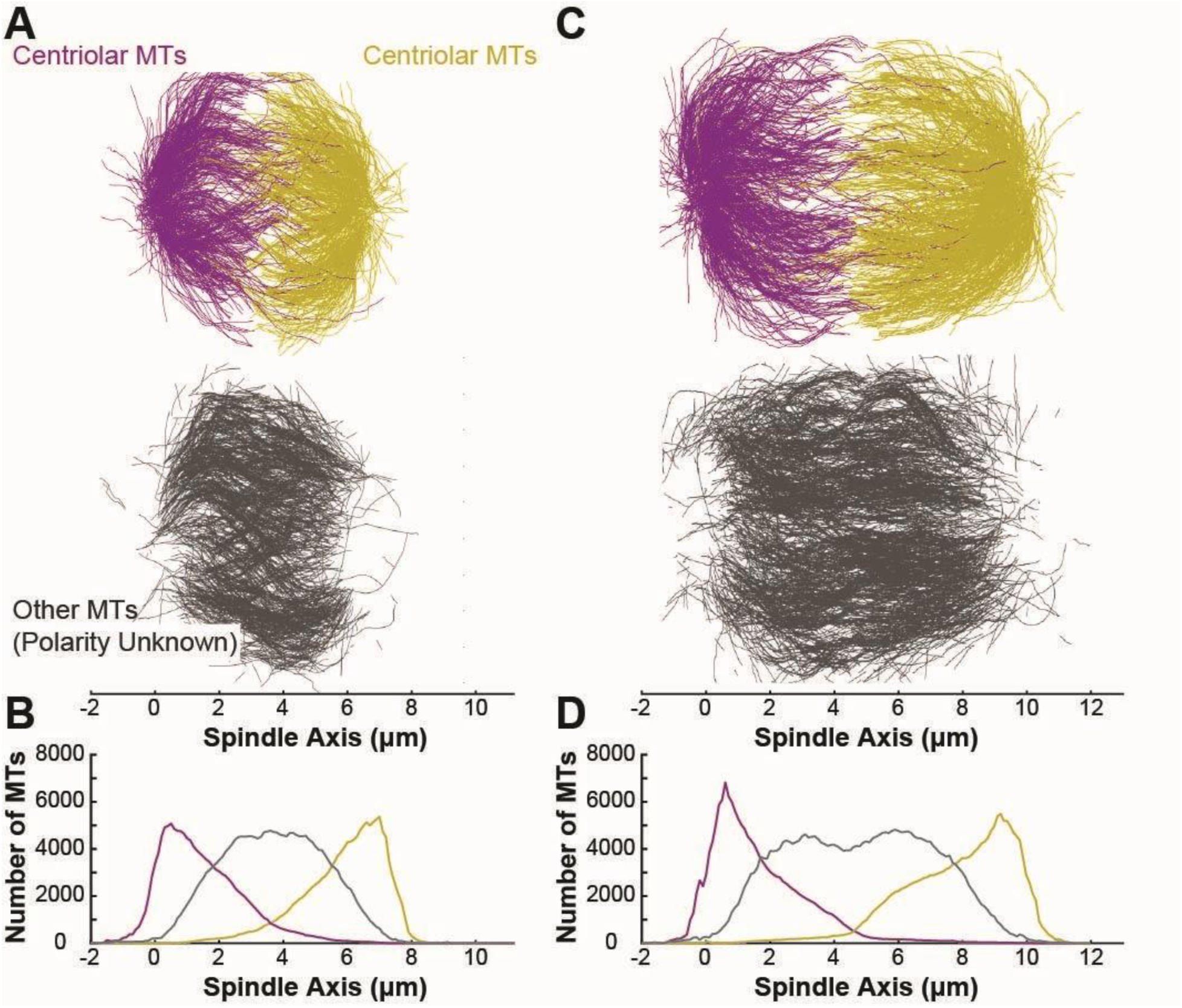
Polarity assignment in resin-embedded serial-section electron tomography reconstructions of HeLa spindles. A) Reconstruction of microtubule trajectories from serial-section, resin-embedded electron tomography of a metaphase HeLa spindle. Microtubules are categorized by distance from either pole. Top: microtubules within 2 µm of the left (purple) or right (yellow) pole. Bottom: microtubules >2 µm from either pole (gray). B) Density of microtubules within 2 µm of the left pole (purple), within 2 µm of the right pole (yellow), and more than 2 µm away from either pole (gray).

**Figure 2 - supplement 1.**
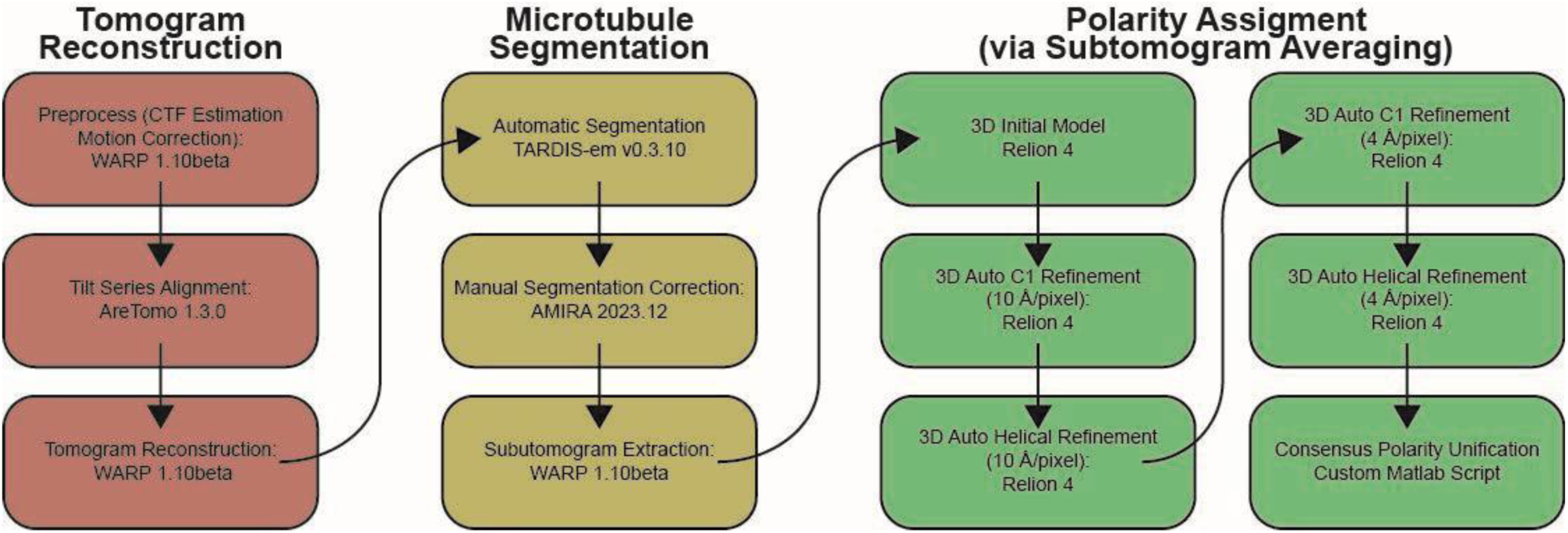
Microtubule polarity assignment pipeline.

**Figure 2 - supplement 2:**
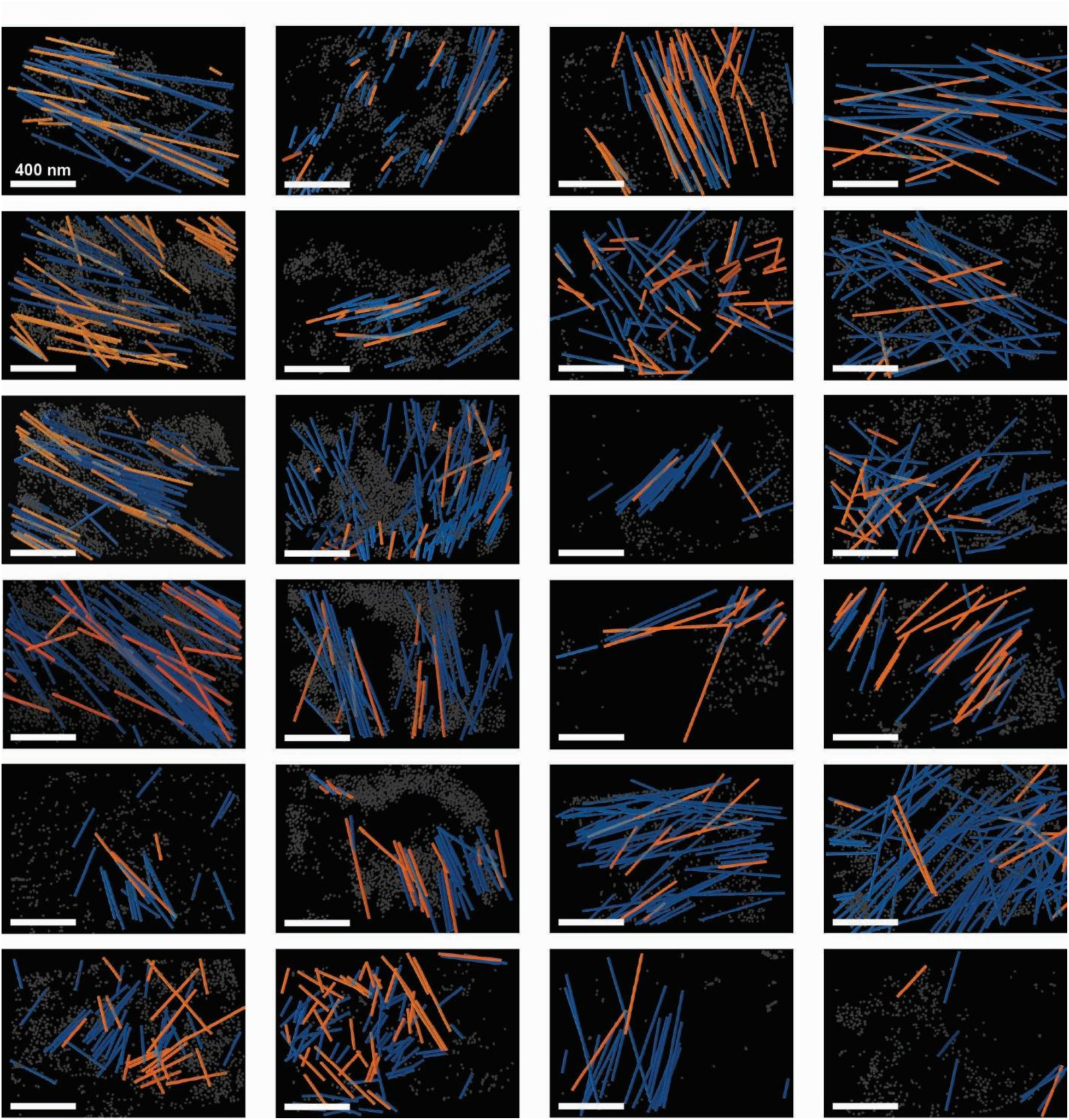
Tomographic reconstructions of metaphase spindles. Blue/orange: microtubules colored by polarity. Gray: ribosomes.

**Figure 3 - supplement 1:**
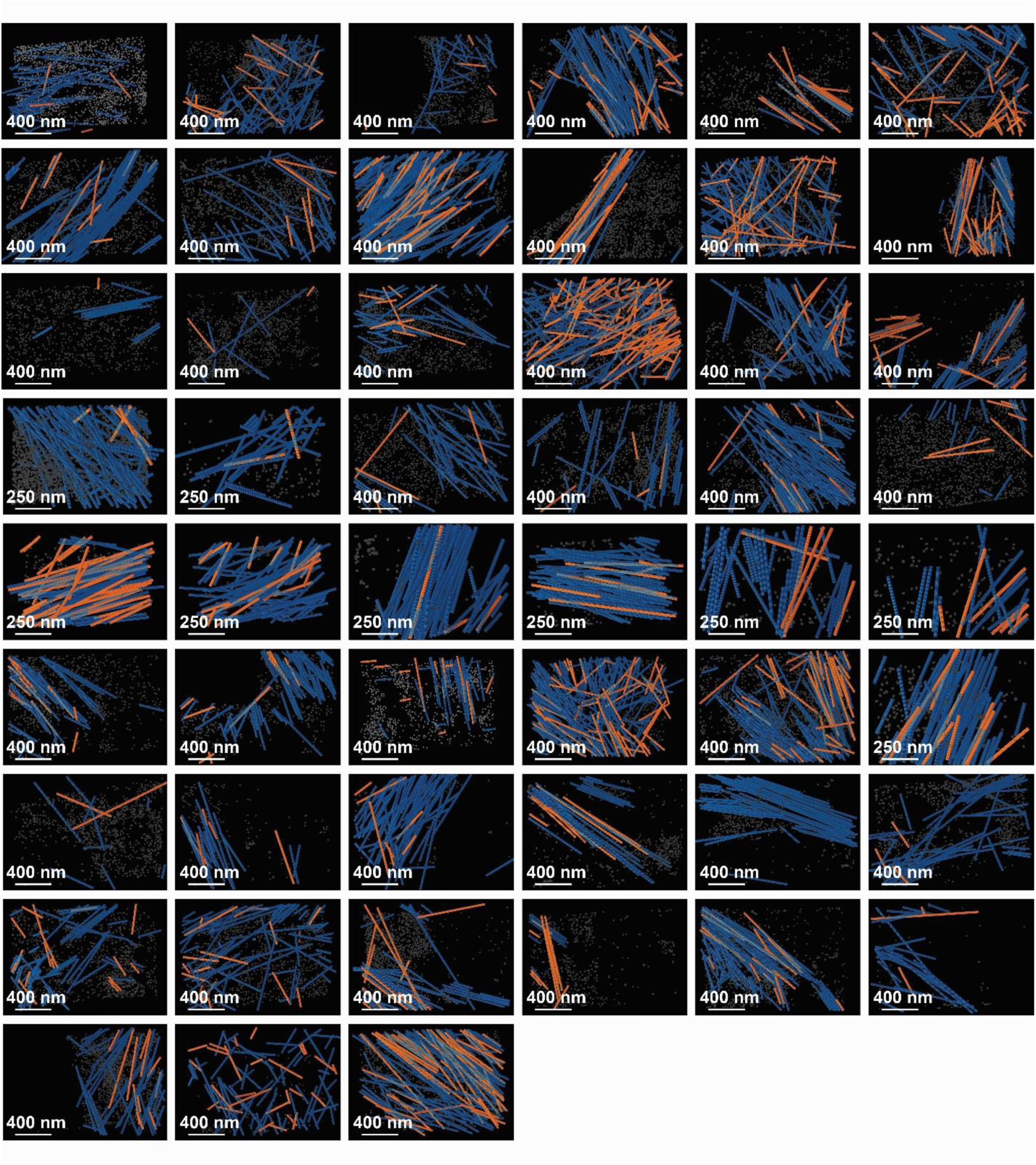
Tomographic reconstructions of Centrinone-treated motor-active monopolar spindles. Blue/orange: microtubules colored by polarity. Gray: ribosomes.

**Figure 4 - supplement 1:**
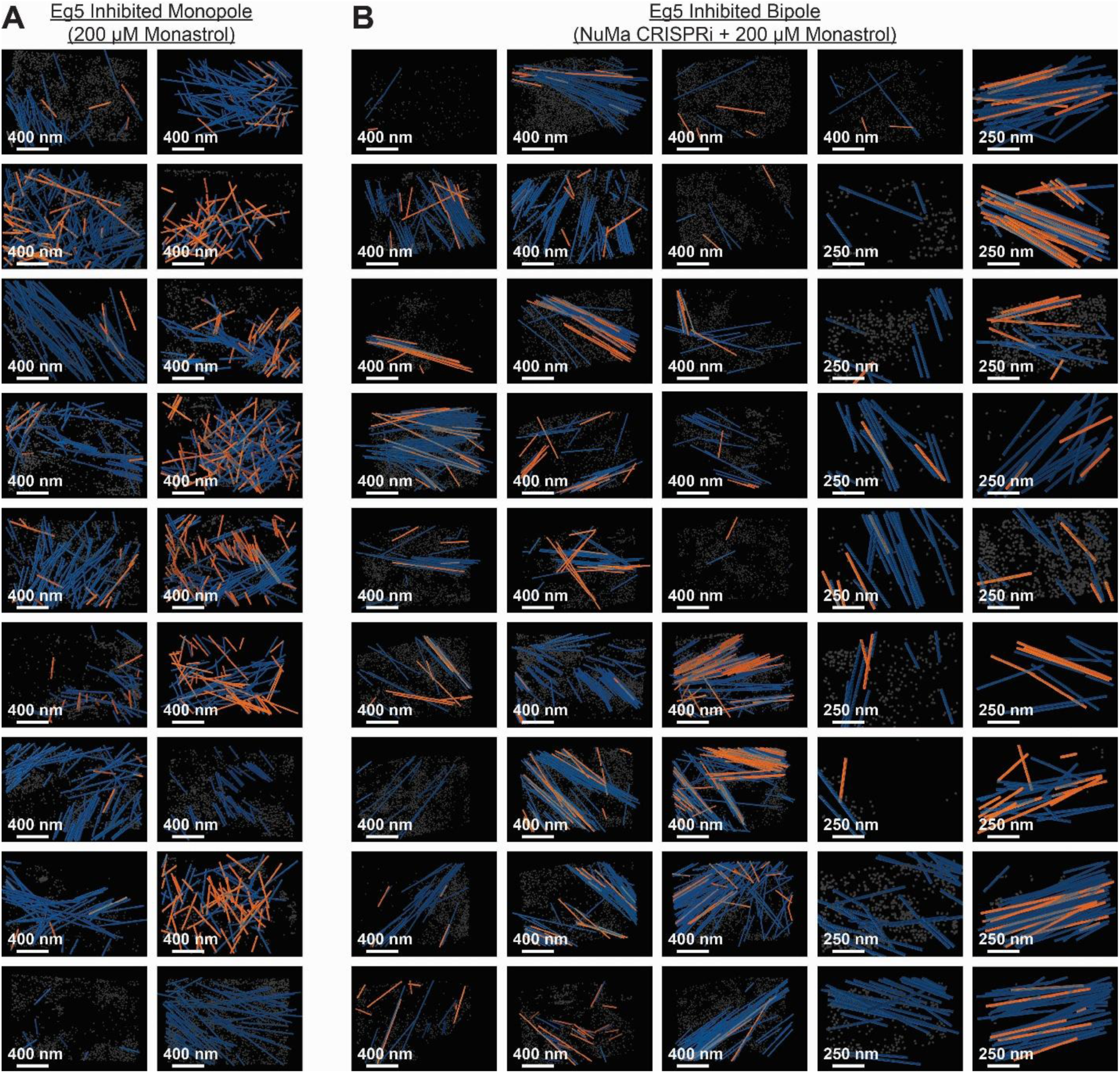
Tomographic reconstructions of Eg5-inhibited monopolar and bipolar spindles. Blue/orange: microtubules colored by polarity. Gray: ribosomes. A) Eg5-inhibited monopolar spindles B) Eg-5 inhibited, NuMa CRISPRi DualKO bipolar spindles

**Figure 4 - supplement 2:**
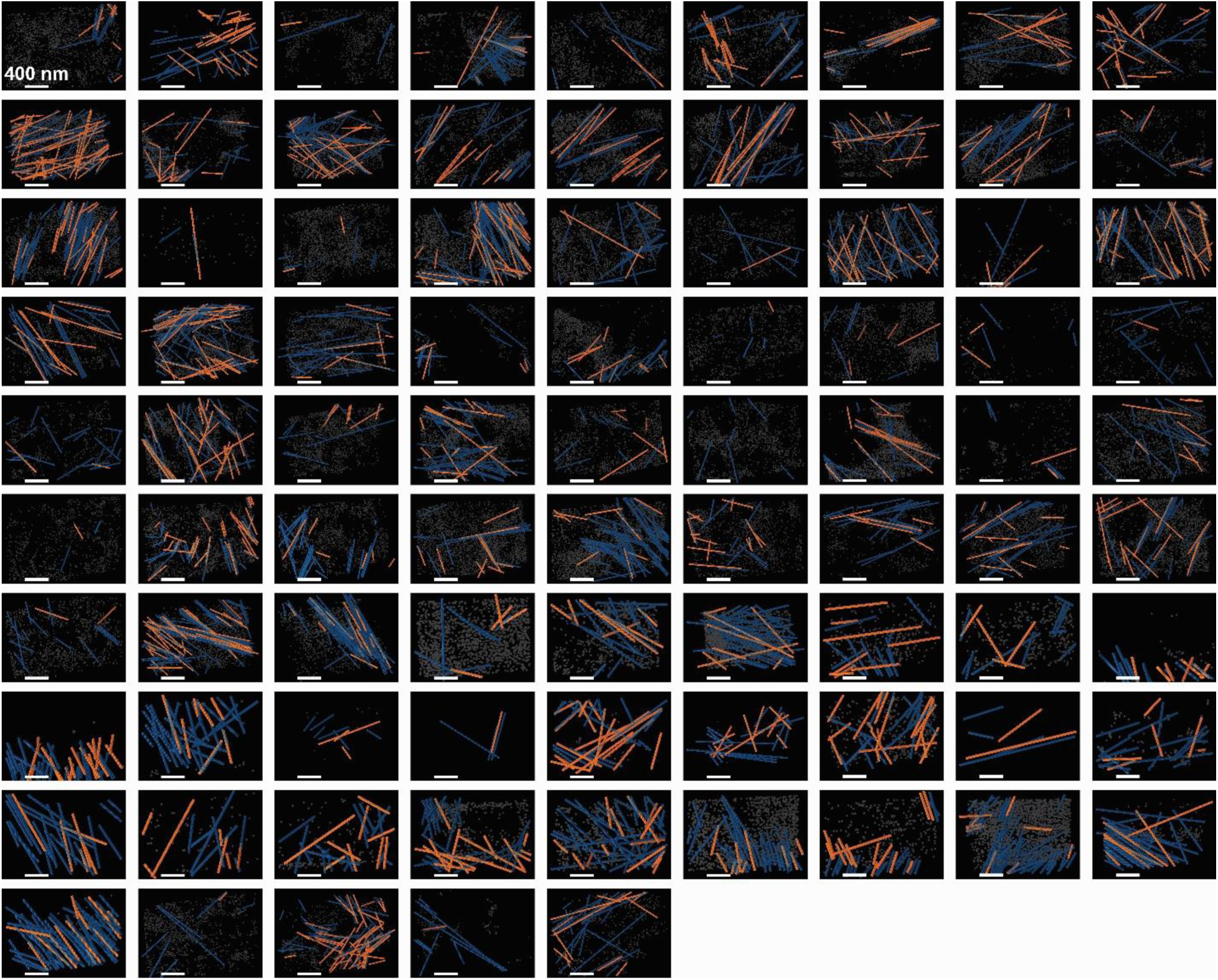
Tomographic reconstructions of Eg5-active NuMa CRISPRi dynein-inhibited spindles. Blue/orange: microtubules colored by polarity. Gray: ribosomes. A) Eg5-inhibited monopolar spindles B) Eg-5 inhibited, NuMa CRISPRi DualKO bipolar spindles

**Figure 6 - supplement 1:**
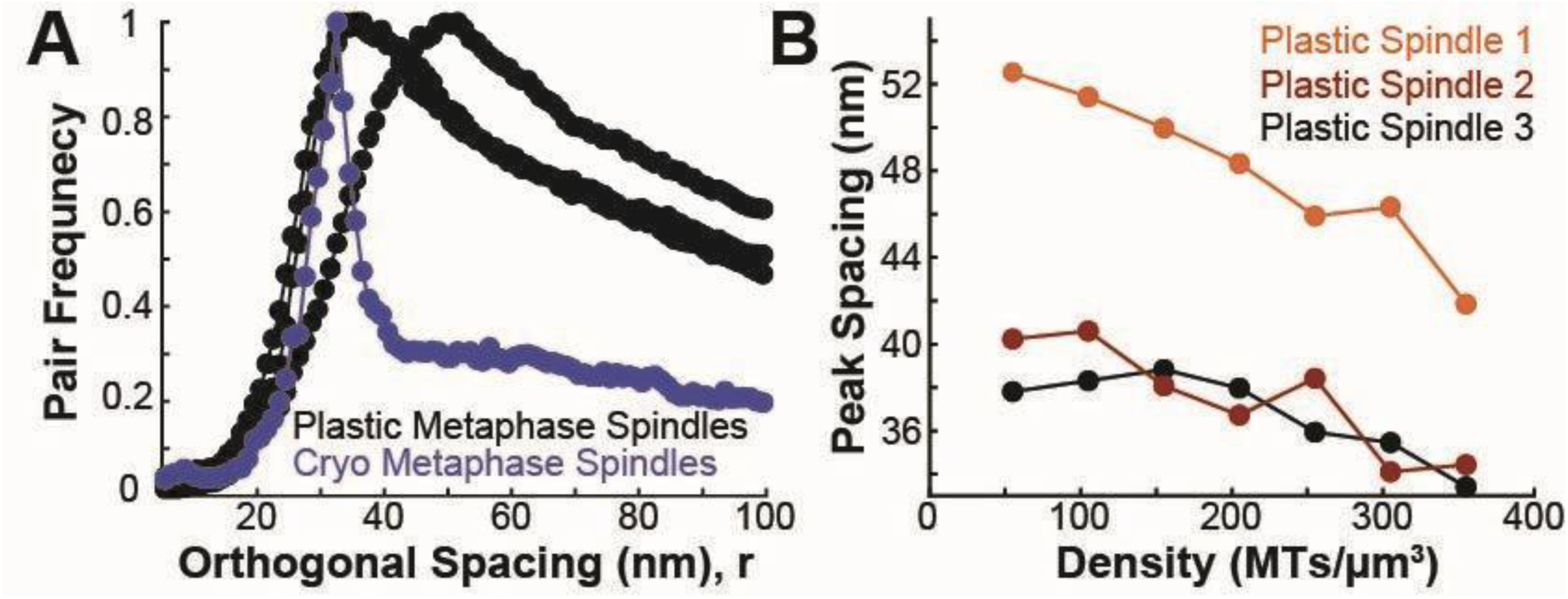
Microtubule density sets spacing in each of the three plastic HeLa spindles. A) Frequency of spacing between microtubules in each of the three plastic spindle reconstructions (black) and cryotomograms (blue). B) Peak spacing between microtubules vs local density for each of the three plastic spindle reconstructions.

